# Silicon-rhodamine functionalized evocalcet probes (EvoSiR) potently and selectively label calcium sensing receptors (CaSR) *in vitro*, *in vivo* and *ex vivo*

**DOI:** 10.1101/2024.02.22.581561

**Authors:** Daniel Bátora, Jérôme P. Fischer, Reto M. Kaderli, Máté Varga, Martin Lochner, Jürg Gertsch

## Abstract

The calcium sensing receptor (CaSR) is a ubiquitously expressed G-protein coupled receptor (GPCR) that regulates extracellular calcium signals via the parathyroid glands. CaSR has recently also been implicated in non-calcitropic pathophysiologies like asthma, gut inflammation and cancer. To date, molecular tools that enable the bioimaging of CaSR in tissues are lacking. Based on *in silico* analyses of available structure-activity relationship data on CaSR ligands, we designed and prepared silicon-rhodamine (SiR) conjugates of the clinically approved drug evocalcet. The new probes EvoSiR4 and EvoSiR6, with differing linker lengths at the evocalcet carboxyl end, both showed a 6-fold and 3-fold increase in potency towards CaSR (EC_50_<45 nM) compared to evocalcet and the evocalcet-linker conjugate, respectively, in a FLIPR^®^-based cellular functional assay. The specificity of the EvoSiR probes towards CaSR binding and the impact of albumin was evaluated in live cell experiments. Both probes showed strong albumin binding, which facilitated the clearance of nonspecific binding interactions. Accordingly, in zebrafish embryos, EvoSiR4 specifically labelled the high CaSR expressing neuromasts of the lateral line *in vivo*. EvoSiR4 was also assessed in human parathyroid tissues *ex vivo*, showing a specific absolute CaSR associated fluorescence compared to parathyroid autofluorescence. In summary, functionalization of evocalcet by SiR led to the preparation of potent and specific fluorescent CaSR probes. EvoSiR4 is a versatile small molecular probe that can be employed in CaSR-related biomedical analyses where antibodies are not applicable.

## Intoduction

Beyond its established role as an intracellular second messenger, calcium (Ca^2+^) is also recognized as an important extracellular signaling molecule^1,2^. The calcium sensing receptor (CaSR) is the major class C G-protein coupled receptor (GPCR) that integrates extracellular calcium signals, and controls calcium homeostasis in the parathyroid glands^3^ and kidneys^4^ by regulating parathyroid hormone production. However, the apparently ubiquitous expression of the CaSR across tissues^5^ underscores the less understood roles of CaSR signaling in a wide variety of physiological processes, for instance its involvement in airway smooth muscle contractility^6^, inflammation^7^, taste modulation^8^ and neuronal excitability^9,10^. Furthermore, dysfunctional CaSR signaling has been implicated in non-calcitropic diseases such as asthma^11^, colorectal inflammatory conditions such as inflammatory bowel disease (IBD)^12^ as well as Alzheimer’s^13^ and cardiovascular disease (CVD)^14^, making this GPCR an attractive potential target for the development of novel therapeutic interventions and as possible disease marker. The recent surge in research on the possible roles of CaSR and its physiological roles therefore requires the development of versatile molecular tools to visualize, monitor and modulate CaSR. While numerous diverse ligands have been developed to modulate CaSR activity^15^, imaging probes remain scarce. Given the limitations of using antibodies, small molecule probes offer versatile biomolecular tools for life imaging^16^.

To date, the development of the small molecular chemical space for CaSR binding ligands has been largely spurred by clinical needs in the therapeutic areas of hyperparathyroidism and osteoporosis^17^. It comprises of L-amino acid and peptide-based moieties that act as positive allosteric modulators (PAMs) at the extracellular Venus-flytrap (VFT) domain and arylalkylamine or quinazolinone containing PAMs and negative allosteric modulators (NAMs) that are specific to the seven transmembrane (7TM) domain of the receptor^18^. To our knowledge, empirical data on the impact of larger fluorophore conjugates such as found in fluorescent small molecular conjugates (FMSCs) on CaSR binding potency and specificity are lacking.

FMSCs have been extensively used for the exploration of ligand binding sites, the quantification of ligand receptor binding interactions in competition experiments, the visualization of receptor expression in cells and tissues, and in providing a fluorescent readout of target engagement^19^. The major objectives in the optimization of FMSCs starts with the selection of the appropriate chemical scaffold to which the introduction of large fluorescent tags does not diminish protein binding affinity. Interrogation of target specificity is a critical, which depends on inherent features of the chemical scaffold, and the optimization of both the linker and the fluorophore. High fluorogenicity of the probe in a receptor-bound state is favorable. High albumin binding can further reduce non-specific background signals in vascularized tissues and enhance the signal-to-noise ratio when washing steps during the labeling procedure are feasible.

In the current study, we generated the first CaSR-binding FMSCs called EvoSiR, which were designed from the molecular scaffold of evocalcet, an FDA-approved PAM calcimimetic drug^20^. Based on SAR analyses, the evocalcet carboxyl group was found to be optimal for SiR functionalization, which was also confirmed by two recent Cryo-EM studies^21,22^. Using a validated FLIPR^®^-based functional CaSR assay, spectrofluorometric analysis, as well as *in vitro, ex vivo* and *in vivo* CaSR labeling studies, we demonstrate that EvoSiR is potent and specific enough to be applied for live imaging, even without extensive washing with albumin.

## Materials and Methods

### Chemicals

Calcein-AM (cat. no.: 22003) and Fluo-8 AM (cat. no.: 21080) were purchased from AAT Bioquest, aliquoted to 50 µg and stored at −20 °C. 4-(4-Diethylaminostyryl-1-methyl-pyridinium-iodid (DiAsp) and NPS-2143 hydrochloride were purchased from Sigma-Aldrich and were kept at −20 °C at a stock concentration of 50 mM in DMSO.

### General procedures for chemical synthesis

The chemical synthesis of EvoSiR probes are depicted in the results section. The detailed description of all synthesis procedures plus the ^1^H NMR, ^13^C NMR and HRMS data are reported in the supplementary material. The purity of the molecules was also assessed using HPLC. All reactions requiring anhydrous conditions were performed in heat-gun, oven or flame dried glassware under inert atmosphere (N_2_ or Ar). Silica gel 60 Å (40–63 mm) from Sigma-Aldrich was used for dry loads. Flash column chromatography was performed on a Teledyne Isco CombiFlash® Rf+ with the corresponding RediSep® prepacked silica cartouches unless otherwise stated. Thin layer chromatography (TLC) was performed on Machery & Nagel Alugram® xtra SIL G/UV 254 visualization under UV light (254 nm) and/or (366 nm) and/or by dipping in anisaldehyde stain and subsequent heating.Commercial reagents and solvents (Acros Organics, Fluorochem, Grogg Chemie, Hänseler, Sigma-Aldrich, Lubio) were used without further purification unless otherwise stated. Dry solvents for reactions were distilled and filtered over columns of dry neutral aluminium oxide under positive argon pressure. Solvents for extraction and flash chromatography were used without prior purification.

^1^H and ^13^C NMR spectra were recorded on a Bruker AVANCE-300 or 400 spectrometers operating at 300 or 400 MHz for ^1^H and 75 or 101 MHz for ^13^C at room temperature unless otherwise stated. Chemical shifts (δ) are reported in parts per million (ppm) relative to tetramethylsilane (TMS) calibrated using residual signals of the solvent or TMS. Coupling constants (J) are reported in Hz. HRMS analyses and accurate mass determinations were performed on a Thermo Scientific LTQ Orbitrap XL mass spectrometer using ESI ionisation and positive or negative mode. HPLCs were measured on a Thermo-Scientific UltiMate 3000 HPLC with H_2_O + 0.1% TFA and MeCN + 0.1 % TFA as eluents on an Acclaim ™ 120 C18 5 µm 120 Å (4.6 x 150 mm) column. The silica-rhodamine carboxylic acid (SiR-CO_2_H) and the corresponding N-hydroxysuccinimide ester (SiR-NHS) was prepared according to procedures previously described elsewhere^23^.

### *In silico* analysis of chemical scaffolds

Assays with a measured CaSR activity (AID) of at least 20 entries were queried from PubChem. Chemical structures were represented in the PubChem 2D fingerprint format, which encodes the presence or absence of chemical substructures in a compound using 881 bits^24^. Clustering was performed using the k-means algorithm^25^, and the optimal number of clusters were selected by applying the *kneedle* algorithm on the inertia scores^26^. Within each AID, we calculated the Tanimoto similarity index for each molecule and the most potent molecule of the assay^27^. For the quadrant analysis, a cutoff value of 0.1 µM was used for the CaSR activity concentrations (AC_50_), below which a compound was considered as a potent CaSR ligand.

### Functional CaSR assay, cell culture

Hamster lung fibroblasts (CCL39) expressing the recombinant human CaSR (HCAR) and non-transected controls were kindly provided by Novartis AG and were kept in standard culturing conditions (37 °C, 5% CO_2_) in ½ DMEM (Roth, 9005.1) – ½ F-12 (Sigma, N4888-500ml) media containing 10% FBS. Media for the HCAR cells contained 500 µg/ml Geneticin™ as a selection antibiotic (ThermoFisher, 10131035). Upon reaching 90-95% confluence, cells were detached and seeded into 96 well-plates (PerkinElmer, 6005182) at a concentration of 35,000 cells per well and were incubated for 24 hours before the experiment. For the functional CaSR calcium assay, cells were first washed twice with HBSS medium (Sigma, H6648), and were incubated in HBSS containing 5 µM Fluo-8^®^ AM and 0.1 mM CaCl_2_ for 30 minutes at 37 °C. Calcium transients were measured upon addition of 0.5 mM CaCl_2_ with the FLIPR Tetra^®^ high-throughput cellular screening systems using the 470 – 495 nm LED excitation and 515 – 575 nm bandpass emission filters. Probe solutions contained 0.5% DMSO and were added to the cells 10 minutes prior to the measurement. dF/F_0_ values were calculated as a change in the peak fluorescence value of the transient for the first 200 data points normalized to the background fluorescence of the well.

### Photophysical characterization of EvoSiR

The photophysical characterization was carried out using the Agilent Cary Eclipse spectrofluorometer and the Agilent Cary 60 UV-VIS spectrophotometer. For the calculation of the extinction coefficient (ε), absorbance was measured for the concentrations of 150-1350 nM for all compounds and the cuvette length was 1 cm. For relative quantum yield measurements (Φ), the Φ_r_ value of 0.68 for the well-characterized Rhodamine B was taken in 94% ethanol as a reference standard^28^. For Rhodamine B, peak absorbance values at 545 nm were compared to the λ_em_ of 567 nm. For the SiR probes, peak absorbance values at 654 nm were compared to the λ_em_ of 670 nm. All measurements were carried out in 94% ethanol at 750nM concentration of the compound. The Φ for each compound was calculated using the following equation:

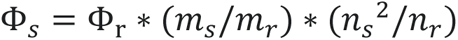

where Φ_s_ depicts the quantum yield of the sample, Φ_r_ corresponds to the quantum yield of the reference solution, m_s_ corresponds to the slope of the absorbance-fluorescence regression plot of the sample, m_r_ depicts to the slope of the absorbance-fluorescence regression plot of the reference solution, and n_s_, n_r_ correspond to the refractive index of the sample and reference solution solvent, respectively, which was 1.3617 for 94% ethanol. For the determination of fluorescence lifetimes (τ), the scan gate time was 0.005 ms, delay time was 0 ms, the number of flashes was 10, the total decay time was 1 ms and the decay curves were determined by averaging 20 cycles.

### Spectrofluorometric analysis of fluorogenicity and quenching

Spectrofluorometric measurements were carried out using the Agilent Cary Eclipse instrument. The voltage of the photomultiplier tube (PMT) was set to 600 for EvoSiR4 and EvoSiR6, and for 500 for SiR (**4**). Fluorescence of the probes were measured at concentrations ranging from 30-3000 nM in PBS containing various amounts of bovine serum albumin (BSA) (Sigma, A7906). Quenching experiments for the calculation of BSA binding were performed by adding 1:1 molar ratio of tannic acid dissolved in _dd_H_2_O (Sigma, 403040) to the solution prior to measuring fluorescence.

### *In vitro* labeling of stably transfected CaSR expressing and non-transfected control cells

HCAR (CaSR+) and CCL39 (CaSR-) cells were kindly obtained from Novartis AG (Basel, Switzerland) and validated in house. Cells were seeded into µ-slide 18 well plates (ibidi^®^, 81816) at a density of 5,000 cells per well and were incubated for 24 hours under standard culturing conditions. As our preliminary data indicated considerable non-specific labeling of dying cells and cellular debris, a viability staining was introduced to ensure probe specificity^29^. Thus, after washing twice with HBSS, cells were incubated in 10 µM Calcein AM solution containing 0.02% Pluronic^®^ F-127 and 0.1% DMSO for 45 minutes at 37 °C. The solution was then removed, washed twice in HBSS before the addition of the HBSS solution containing the SiR probes. All probe solutions were protected from light exposure and contained 0.5% DMSO and various concentrations of BSA depending on the experiment. Incubation with the fluorescent probes was 20 minutes long, after which a single washing step was introduced with HBSS before imaging. Images of the cells (Calcein: FITC channel, SiR: Cy5 channel) were acquired with a Nikon Eclipse TI2 inverted widefield fluorescence microscope using a 60x magnification oil immersion objective (Nikon CFI Plan Fluor, 60x, NA=0.85). For each well, a minimum of 3 images were acquired. For each image, cells were segmented to generate labels on the FITC channel post-acquisition using the deep neural network based Cellpose package in Python 3.9 (model_type: cyto2, diameter: 200, flow_threshold: 0.4)^30^. To normalize pixel intensity values across images, for each image, a background value was calculated by taking the average pixel intensity value of the pixels in which no cellular label was present and were negative for the EvoSiR label (triangle thresholding used). For each cell, the skewness of the value distribution, and the mean of the Cy5 pixel intensities were calculated.

### Zebrafish husbandry

Transgenic *Tol056*^31^ and *Tg(7xTCF-Xla.Siam:GFP)*^32^ zebrafish (*Danio rerio*) used in this study were maintained and bred within the fish facility of ELTE Eötvös Loránd University (Budapest, Hungary) according to standard protocols^33,34^. The protocols used in this study were approved by the Hungarian National Food Chain Safety Office (Permit Number: XIV-I-001/515-4/2012).

### In vivo labeling and imaging of zebrafish embryos

For the neuromast labeling experiments, wild-type 4-6 days post fertilization (dpf) zebrafish larvae were incubated in 200 µl E3 medium containing different doses of the SiR probes (500 – 5000 nM), 2.5 µM of 4-(4-Diethylaminostyryl-1-methyl-pyridinium-iodid (DiAsp, Sigma, D3418), which specifically labels hair cells in the lateral line in 0.5 % DMSO for 30 minutes^35^. Next, washing and anesthesia was achieved by placing the larvae in a 35 mm imaging dish (MolBiTec, Imaging dish 1.5) which contained 500 µl E3 medium with 168 mg/l MS-222 (Tricaine Mehanesulfonate, Sigma) for 10 minutes. After removing the solution, fish were side-embedded in the imaging dish using 500 µl 0.75 % low-melting temperature agarose (Sigma, A4018). Widefield imaging was done using the Zeiss Axio Observer fluorescence microscope (LD A-Plan 10x objective, NA=0.25), confocal imaging was done using the Zeiss LSM 800 confocal fluorescence microscope (LD LCI Plan-Apochromat 25x objective, NA=0.8).

### Zebrafish startle reflex analysis

For the behavioral analysis, wild-type 4-6 dpf zebrafish larvae were placed individually into custom-built sound delivery system described in a previous publication^36^. For acoustic stimulation, we used a 90 dB stimulus, and recorded the subsequent behavioral responses using a high-speed camera (xiQ USB3 vision) at a framerate of 500 fps using custom scripts written in Python. The camera was placed 30 cm above the sound delivery system. The latency of the startle reflex was quantified manually by using a LED flash (780 nm wavelength) in parallel with the time of stimulus delivery. Escape responses were quantified manually, where we considered a short-latency startle response between 2-16 ms, and a long-latency startle response above 16 ms.

### *Ex vivo* labeling of surgically resected human parathyroid adenomas

The part of the study involving human tissues has been registered at clinicaltirals.gov and can be retrieved under the ID: NCT03831620. The study was approved by the ethics commission of the canton Bern, Switzerland (KEKBE 2018-02218). Written informed consent was obtained from all patients. Human parathyroid tissues were surgically resected and were immediately put on dry ice and stored at −80 °C. The tissues were thawed on 4 °C and were washed with PBS thoroughly. The labelling of the tissues was achieved using 500 nM of EvoSiR4 for 30 minutes in PBS containing 0.1% BSA. After labelling, the tissues were washed twice with PBS containing 0.1% BSA. For evocalcet competition experiments, the tissues were co-incubated with EvoSiR4 and evocalcet in a 1:5 molar ratio for 30 minutes and then washed twice with PBS containing 0.1% BSA. The images were acquired using an IVIS® Spectrum *in vivo* imaging system with a λ_ex_ of 780 nm and λ_em_ of 840 nm for the autofluorescence (ICG channel) and with a λ_ex_ of 650 nm and λ_em_ of 670 nm for the EvoSiR label (Cy5 channel).

### Statistics and data analysis

All statistical analysis was carried out in Python 3.9 using the SciPy package. Data have been obtained in at least 3 independent experiments and are represented as mean ± standard deviation unless otherwise indicated. Statistical significance was determined using Mann-Whitney test with the Bonferroni correction where applicable unless otherwise stated (*p<0.05, **p<0.01, ***p<0.001, ****p<0.0001). No statistical tests were applied to predetermine sample sizes, but our sample sizes were similar to those of previous reports.

## Results

### Selection of optimal CaSR ligands for the incorporation of fluorescent moieties

To decipher the structural diversity of CaSR-binding molecular scaffolds, a total of 840 unique molecules were gathered *in silico* from PubChem activity assays (AID) with their empirical active concentration (AC_50_) values towards CaSR. Activity assays that contained at least 20 molecules were selected, which resulted in a total of 556 molecules. The dimensionality of the 881-bit PubChem substructure fingerprints of the molecules were reduced to 20 latent dimensions using principal component analysis (PCA). The first 10 PCs could explain 73% of the variance present in the data (Figure S1). We next calculated the correlation matrix from the 10 PCs and clustered the structures using the *k*-means clustering algorithm. The optimal number of clusters was eight, which was calculated from the inertia values (Figure S1). The chemical structures of the molecules closest to the cluster centers are represented in Figure 1. Next, we assessed which molecular scaffolds are most tolerant towards structural modifications without losing potency. To this end, within each AID, the AC_50_ value of each molecule was plotted as a function of the Tanimoto similarity value to the most potent molecule (Figure 1). The number of molecules within the quadrant with dissimilar structures and low AC_50_ values (Q_3_) normalized to the molecules in the quadrant with dissimilar structures and high AC_50_ values (Q_1_) and similar structures and high AC_50_ values (Q_2_) were quantified. Only 5 AIDs (459797, 738057, 744122, 1464365 and 1802224) showed a positive value whereas AID 1464365 had the highest value, suggesting that this scaffold might be tolerant towards the incorporation of fluorescent moieties.

**Figure 1.**
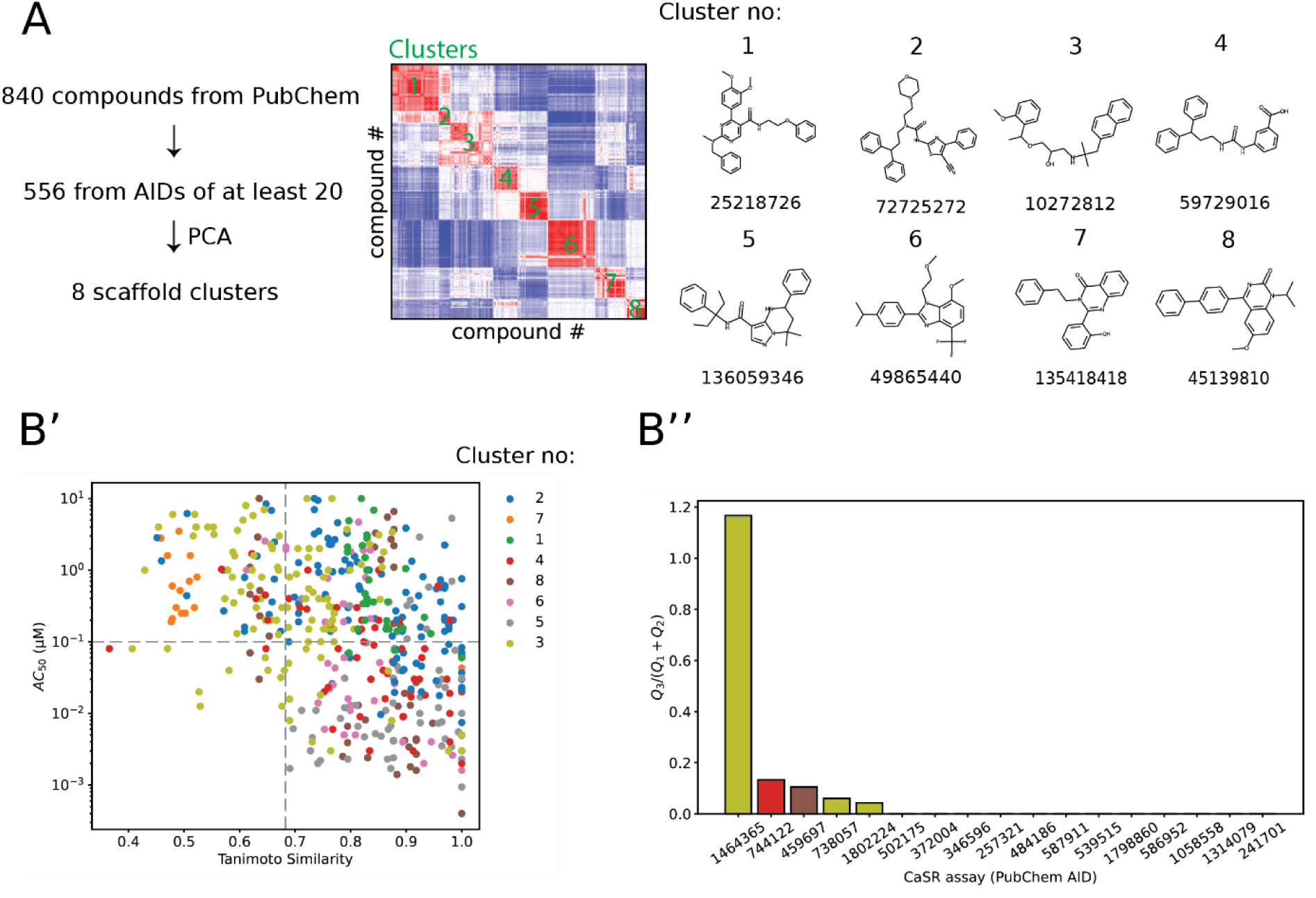
*In silico* analyses reveal optimal chemical scaffolds for the functionalization with fluorophores. (A) Description of the workflow to obtain 2D structure-based clusters on CaSR-binding molecules downloaded from PubChem. The heatmap depicts the correlation matrix of the Tanimoto-similarity of 10 latent dimensions of the 2D fingerprints for all compound, which identified eight chemical clusters (green text). On the right, the seven most representative molecules and their PubChem ID from each chemical cluster is shown (closest Euclidean distance to the coordinates of cluster centroids). (B’) Scatter plot depicting the relationship between the Tanimoto similarity of each molecule to the most potent molecule of each assay (AID) and the half-maximal activity value (AC_50_). (B’’) The ratio between the number of highly potent (<100 nM) and dissimilar molecules (Q_3_) to molecules with reduced potency (>100 nM) and high (Q_1_) or low (Q_2_) similarity is quantified. The assay with the AID 1464355 had the highest number of dissimilar and potent molecules.

### Synthesis of EvoSiR

Based on the *in silico* data and reported cryo-EM structures^21,22^, evocalcet was extended at the carboxyl end with linkers prior to functionalization with SiR (Scheme 1). Using 1-ethyl-3-(3-dimethylaminopropyl)carbodiimide (EDC) in the presence of DMAP efficiently coupled mono-protected bis-amines to deliver the corresponding amides **1** and **2**. Cleavage of the Boc-protecting group of 2 by treatment with TFA gave the free amine **3**, which was reacted with the activated dye ester SiR-NHS to yield the final probe EvoSiR6. For EvoSiR4, we found it more advantageous to first couple the mono-protected bis-amine linker with silicon-rhodamine (SiR-CO_2_H). After Boc-deprotection, the free amine **4** was subsequently reacted with evocalcet with the aid of EDC and DMAP. The relatively low yields of the final probes EvoSiR4 and EvoSiR6 reflect their challenging chromatographic purification rather than the efficiency of the amide coupling steps. The isolation of the final probes may be optimized in future endeavors.

**Scheme 1.**
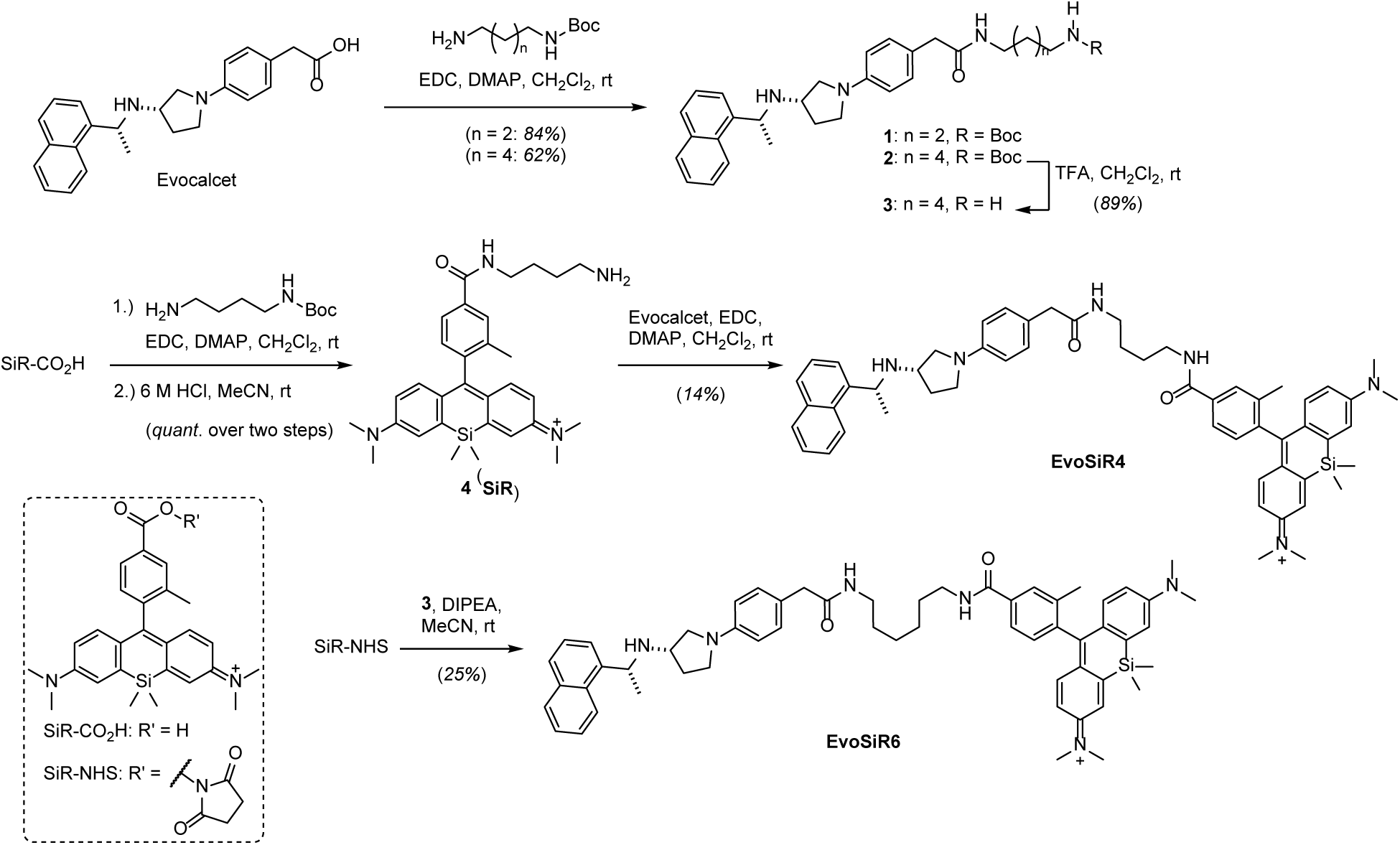
Synthesis of EvoSiR4 and EvoSiR6. Synthesis of the silicon rhodamine conjugate CaSR probes EvoSiR4 and EvoSiR6. Abbreviations: Boc, *tert*-butyloxycarbonyl; EDC, 1-ethyl-3-(3-dimethylaminopropyl)carbodiimide; DMAP, 4-(dimethylamino)pyridine; TFA, trifluoroacetic acid; DIPEA, diisoproplyethylamine; NHS, *N*-hydroxysuccinimide.

### EvoSiR proves have improved CaSR affinity compared to evocalcet in a functional *in vitro* assay

To assess CaSR receptor binding affinity, we quantified the concentration-response of the PAM effect of the synthesized molecules and evocalcet on CaSR activity using an *in vitro* FLIPR^TM^ cellular assay (Figure 2). In our assay, for evocalcet an EC_50_ value of 243±15 nM in response to the addition of 0.5 mM CaCl_2_ was obtained. In comparison, both EvoSiR4 and EvoSir6 exhibited similar but more potent EC_50_ values than evocalcet in the range of 40 nM (Figure 2), whereas the EC_50_ value for evocalcet with only the linker (compound **1**, Scheme 1, Figure S2) was 101±6 nM (Supporting Figure 2). Importantly, the fluorescent property of the EvoSiR probes at the concentrations used did not interfere with the assay fluorescent readout (data not shown). We next modeled the impact of albumin binding on the EC_50_ values for EvoSiR4 by adding different concentrations of bovine serum albumin (BSA). The addition of EvoSiR in 0.1% BSA increased the EC_50_ value to 64±12 nM, 0.5 % BSA to 102±34 nM and 1% BSA to 106±42 nM (Figure 2), indicating saturable BSA binding. Similarly, the hill coefficients of the fitted curves also increased upon addition of BSA. In agreement, washing the molecules incubated in 1% BSA with HBSS upon 10-minute incubation in the cellular assay led to a 10-fold reduction in the EC_50_ value (Figure 2).

**Figure 2.**
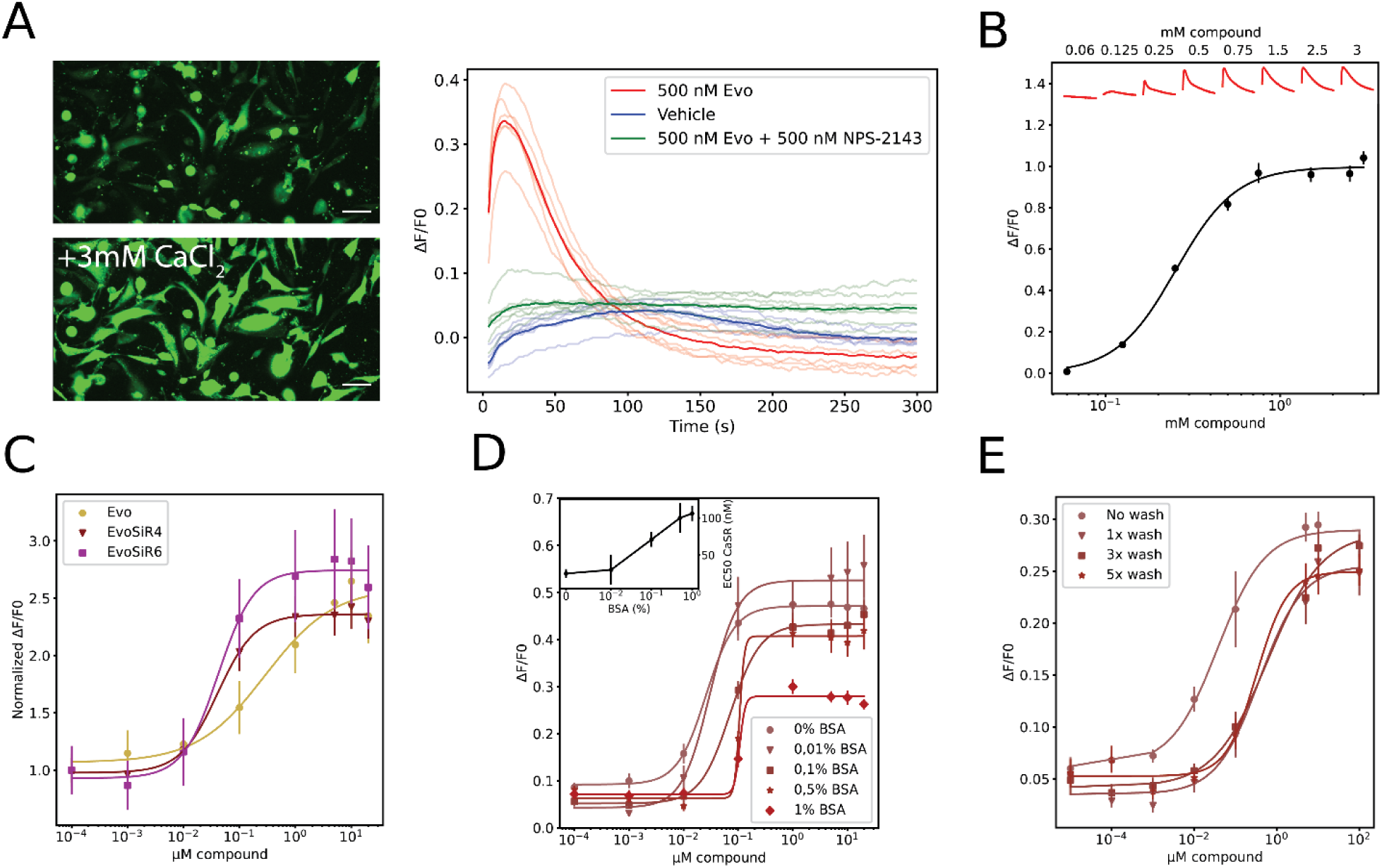
*In vitro* functional CaSR assay reveal retained potency of EvoSiR probes. (A) Left: Example images showing the increase of intracellular calcium in CCL39 cells overexpressing the human CaSR (HCAR) in response to the addition of 3 mM of CaCl_2_ stained with FLuo-8 AM (white scale indicates 15 µm). Right: representative intracellular calcium transients measured by FLIPR in HCAR cells upon the addition of 0.5 mM calcium and either 500 nM of evocalcet (Evo), vehicle and 500 nM evocalcet + the 500nM of the CaSR inhibitor NPS-2143. (B) Concentration-response curve of intracellular calcium release in response to extracellular calcium in HCAR cells. CaSR activity was measured indirectly as a relative increase in fluorescence (dF/F0) (EC_50_=0.252±0.01 mM, n=3, in triplicates). Representative traces for each concentration in mM are shown at the top of the plot in red. (C) Concentration-response curve of evocalcet and the two EvoSiR derivatives (EvoSiR4, EvoSiR6) on the positive allosteric modulation of CaSR activity (n=5-13 in triplicates and EC_50_=0.243±0.015, 0.038±0.009 and 0.042±0.011 µM for evocalcet, EvoSiR4 and EvoSiR6, respectively). (D) The impact of various concentrations of bovine serum albumin (BSA) on the potency of EvoSiR4 on CaSR activity (n=5 each in triplicates for all conditions). The inset indicates the EC_50_ values of each condition as a function of BSA percentage. (E) The impact of washing upon 5-minute incubation HCAR cells with various concentrations of EvoSiR4 and 0.1% BSA (n=3 each in triplicates for all conditions, EC_50_=0.037±0.008 (no wash), 0.38±0.24 (1x wash), 0.62±0.31 (3x wash) and 0.33±0.21 (5x wash).

### Photophysical properties of EvoSiR probes

For SiR (**4**), EvoSiR4 and EvoSiR6, standard photophysical parameters were determined, including extinction coefficients (ε), peak excitation (λ_ex_) and emission (λ_em_) wavelengths (Figure 3), quantum yields (Φ) and fluorescence lifetimes (τ) and are reported in Table 1. Regression curves for the analysis are shown in the supplementary information, Figure S3.

**Figure 3.**
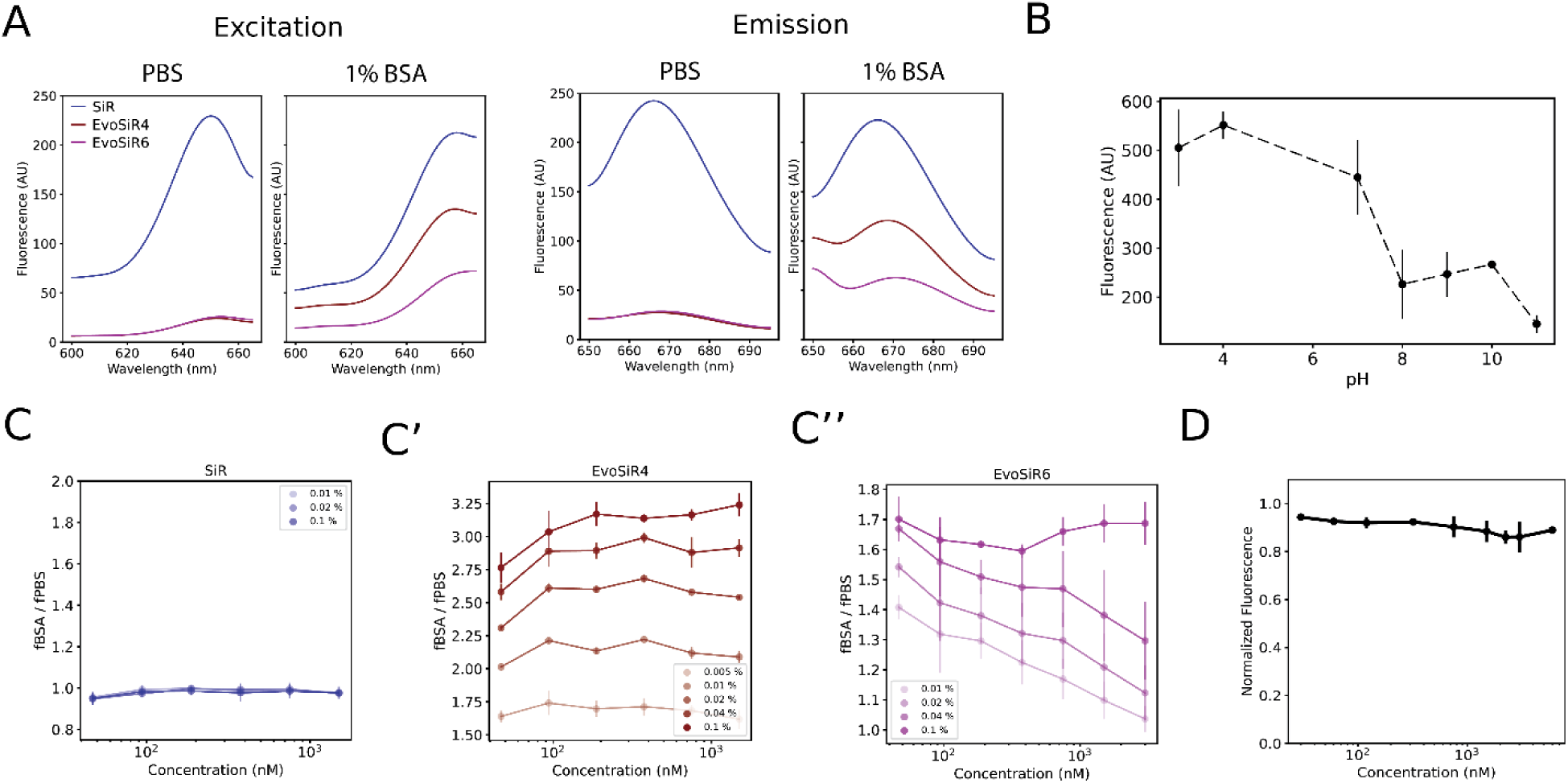
Photophysical properties of EvoSiR and the effect of BSA binding. (A) Plot depicting the fluorescence excitation and emission spectrum of EvoSiR4. The peak excitation wavelength (λ_ex_) was 655 nm, and the peak emission wavelength (λ_em_) was 665 nm. (B) The effect of pH on the peak fluorescence at 670 nm for 750 nM of EvoSiR4 measured in PBS solution (n = 3 each). The absolute fluorescence was highest at below physiological pH, while pH above 10 resulted in a decrease in fluorescence. (C) The effect of BSA binding on the absolute peak fluorescence of SiR (**4**) and EvoSiR probes using various concentrations of BSA. For the analysis, we used 30-1500 nM of SiR (**4**) (C’), EvoSiR4 (C’’) and EvoSiR6 (C’’’) (n=3 each, in triplicates). The values on the y axis represent the absolute fluorescence value in BSA-containing PBS solution normalized to the absolute fluorescence value in PBS. (D) Plot depicting the effect of pre-incubating the BSA-containing PBS solution in various concentrations of evocalcet (30 – 6000 nM) on the fluorescence value of 350 nM of EvoSiR4 (n=6, in triplicates). No significant decrease in the fluorescence was observed, indicating that EvoSiR4 and evocalcet occupy distinct binding sites on BSA.

**Table 1:** Photophysical properties of EvoSiR compounds.

| Compound | $\epsilon(\text{M}^{-1}\text{cm}^{-1})$ | $\lambda_{\text{ex}}(\text{nM})$ | $\lambda_{\text{em}}(\text{nM})$ | $\Phi$ | $\tau(\text{ns})$ |
| --- | --- | --- | --- | --- | --- |
| SiR (4) | 7051 | 650 | 670 | 0.347 | 1.92±0.733 |
| EvoSiR4 | 14175 | 650 | 670 | 0.288 | 2.43±0.164 |
| EvoSiR6 | 16601 | 650 | 670 | 0.278 | 2.33±0.135 |

### Fluorogenicity and BSA binding of EvoSiR probes

We next assessed the fluorogenicity of the probes by measuring the effect of pH and BSA binding by using spectrofluorometry. In PBS, the absolute fluorescence of SiR (**4**) was 10 times higher than of EvoSiR4 and EvoSiR6. The addition of 1% BSA significantly increased the fluorescence of EvoSiR probes, reflecting fluorogenicity upon BSA binding, while it had no effect on SiR (**4**) (Figure 3). We found that the absolute fluorescence of the EvoSiR probes in PBS was strongly dependent on the pH of the solution, with increasing fluorescence values in acidic conditions (Figure 3). In measuring the impact of protein binding on absolute fluorescence, we found no significant change to the absolute fluorescence of SiR (**4**) in response to the addition of BSA in the range of 0.01-0.1% (Figure 3). In contrast, in 0.1 % BSA, EvoSiR4 and EvoSiR6 had a 4-fold and 1.8-fold increased fluorescence compared to PBS, respectively. For EvoSiR4, the BSA-dependent increase in fluorescence was uniform in the concentration range tested (30-1500 nM), whereas for EvoSiR6, this was only observed at 0.1% BSA (Figure 3), suggesting a higher BSA binding ability of EvoSiR4. Next, we pre-incubated the BSA solution with various concentrations of evocalcet. We found no significant difference in the absolute fluorescence between the evocalcet preincubated samples and the control, suggesting that EvoSiR and evocalcet occupy different binding sites in BSA due to the bulky SiR moiety (Figure 3).

### Validation of specificity of EvoSiR in a CaSR overexpressing cell line

*In vitro* labeling of human CaSR stably overexpressing HCAR cells (CaSR+) with EvoSiR4 resulted in highly fluorescent membrane-associated puncta, which were largely absent in the labeling of the non-transfected CCL39 control cells (CaSR-). We therefore used the skewness of the pixel intensity distribution for the evaluation of labeling specificity. For EvoSiR4, significantly higher skewness for HCAR cells was measured at concentrations above 500 nM with no BSA and 0.1% BSA, and for concentrations above 2500 nM with 1% BSA compared to CCL39. Without BSA, the saturation of skewness values appeared at 500 nM for HCAR but was not saturated for CaSR-cells even at 5000 nM (Figure 4). We repeated the same experiment with the EvoSiR6 probe, which similarly differentiated between the CaSR+ and CaSR-cells, confirming the specific labeling (Figure 4). Incubation with SiR (**4**) revealed no significant difference between CaSR+ and CaSR-cells, confirming the specificity of EvoSiR4 CaSR labelling (Figure 4). Assuming identical binding sites of evocalcet and EvoSiR probes and to confirm the labeling specificity, we conducted a competition experiment by pre-incubating the CaSR+ cells with 0-500 nM of evocalcet in 0.1% BSA. Evocalcet concentration-dependently decreases the highly fluorescent puncta, resulting in a significant reduced skewness of the pixel intensity distribution in CaSR overexpressing cells (Figure 4). Washing with BSA showed a divergent effect on the CaSR+ and CaSR-cells, leading in an increased signal background ratio. In the CaSR-cells, washing 5 times with 0.1% BSA resulted in a reduction of the skewness and the normalized mean pixel intensity, reflecting the washout of non-CaSR labelling (i.e. background). In contrast, washing the CaSR+ cells with BSA had no effect on the skewness, but significantly increased the normalized mean pixel intensity (Figure 4). Next, we serendipitously discovered that tannic acid is a highly efficient molecular quencher of the fluorescence of EvoSiR probes and SiR (**4**) with IC_50_ values in the low nanomolar ranges (Figure 4). We therefore tested whether tannic acid can be used as a contrast-enhancing agent for EvoSiR labelling. As shown in Figure 4, in CaSR+ cells labelled with 500 nM of EvoSiR4, the administration of 100 nM of tannic acid indeed resulted in a 40% increase in the signal-to-noise ratio (Figure 4), suggesting that tannic acid quenches the CaSR bound EvoSiR4 less efficiently.

**Figure 4.**
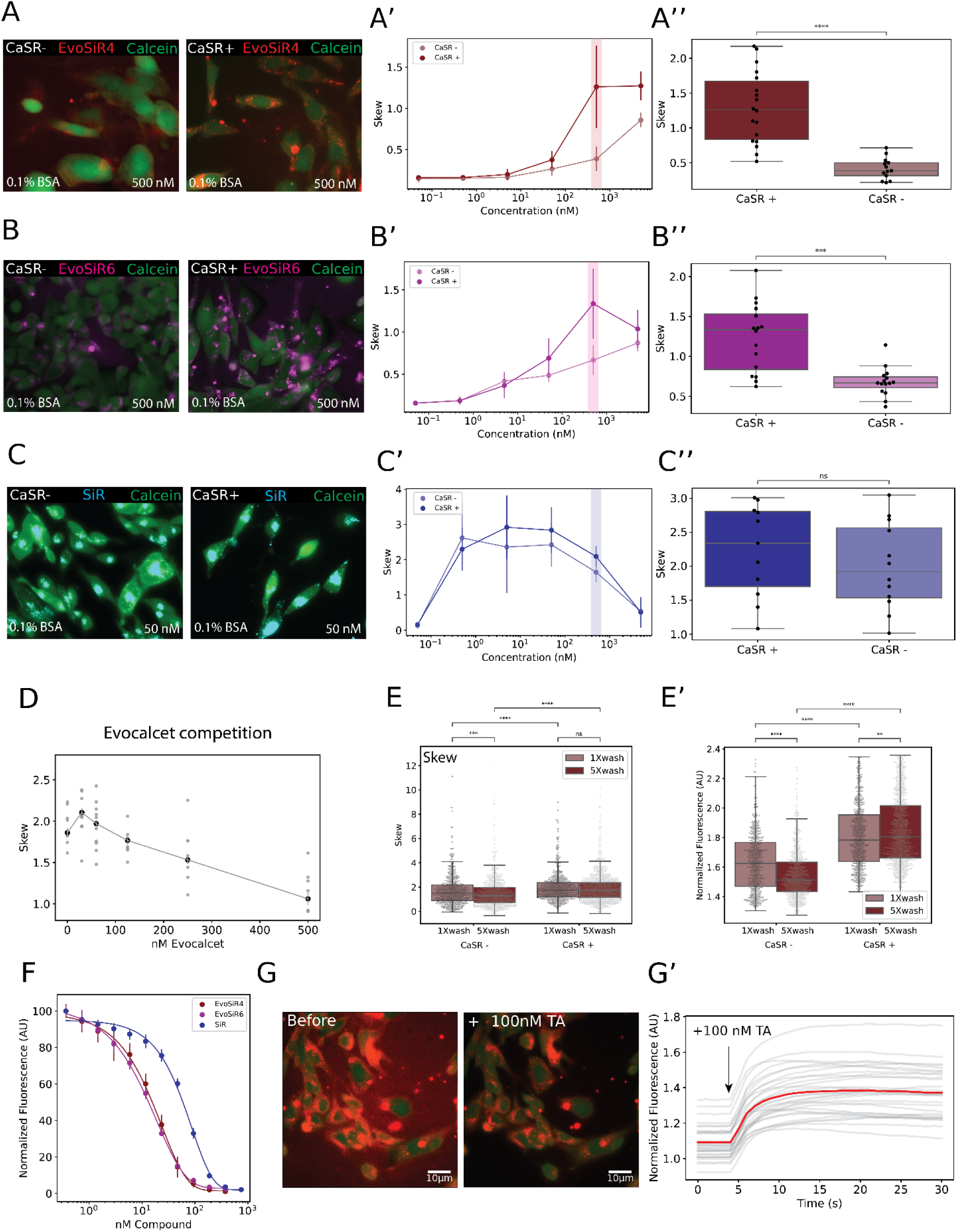
EvoSiR probes specifically label CaSR-expressing cells. (A) Representative images of labelling with 500 nM of the probe of control (CaSR-) and CaSR-expressing cells (CaSR+). Viable cells were identified by staining with Calcein AM. The middle plots depict the concentration-response effect measuring the skewness of the pixel intensity distribution for each cell (at least n = 9-16 independent experiments for each condition). The right plot shows the skew value for each compound at the 500 nM condition (p<0.0001, p=0.0003 0.37, for EvoSiR4, EvoSiR6 and SiR (**4**), respectively; Mann-Whitney Wilcoxon). (D) Pre-incubating the CaSR+ cells with various concentrations of the non-fluorescent CaSR positive allosteric modulator, Evocalcet, which resulted in a concentration-dependent reduction of the EvoSiR4 labelling (n=6 for all conditions). (E) Boxplots depicting the differential effects of washout with 0.1% BSA on the CaSR- and CaSR+ cells. The left plot depicts the skewness of each cell (n =17-25; p=0.99 for the comparison of CaSR+ 1X wash and CaSR + 5X wash, p=0.00012 for the comparison of CaSR-1X wash and CaSR-5X wash, p<0.0001 for all other comparisons, Mann-Whitney Wilcoxon test with Bonferroni correction), whereas the right plot depicts the average normalized pixel intensity (n =17-25; p=0.008 for the comparison of CaSR+ 1X wash and CaSR + 5X wash, p<0.0001 for all other comparisons, Mann-Whitney Wilcoxon test with Bonferroni correction). (F) Concentration-response curve of the fluorescence quenching for 750 nM of SiR probes by the administration of tannic acid measured with a spectrofluorometer. The EC_50_ values are 15.59±4.32 nM, 14.95±2.65 and 58.17±4.86 for EvoSiR4, EvoSiR6 and SiR (**4**), respectively (n=3, in triplicates). (G) On the left, the two representative images show the effect of administration of 100 nM of tannic acid on the signal-to-noise ratio of EvoSiR4 labelling on CaSR+ cells. On the right, the plot depicts the 40% increase in signal-to-noise upon the addition of 100 nM of tannic acid in a timelapse imaging experiment, where the red line indicates the mean signal-to-noise ratio, and the gray line indicates each cell.

### EvoSiR4 and EvoSiR6 specifically label hair cells of neuromasts of the zebrafish lateral line organ *in vivo*

To confirm the specificity of EvoSiR probes in an *in vivo* setting, we next assessed labeling specificity of hair cells of the neuromasts within the lateral line organ of 4-6 days post fertilization (dpf) larval zebrafish, which has been previously confirmed to highly express CaSR^37^. As expected, upon addition of either EvoSiR4 or EvoSiR6, we observed a marked increase in the fluorescence intensity within the neuromasts, whose identify we confirmed pharmacologically by co-labeling with the hair cell specific dye DiAsp^35^ (Figure 5). This increase in fluorescence for the EvoSiR probes was significantly higher compared to SiR (**4**), which also labeled other cells and exhibited a markedly higher background fluorescence. In agreement with their comparable CaSR EC_50_ values, no significant difference between EvoSiR4 and EvoSiR6 was observed (Figure 5).

**Figure 5.**
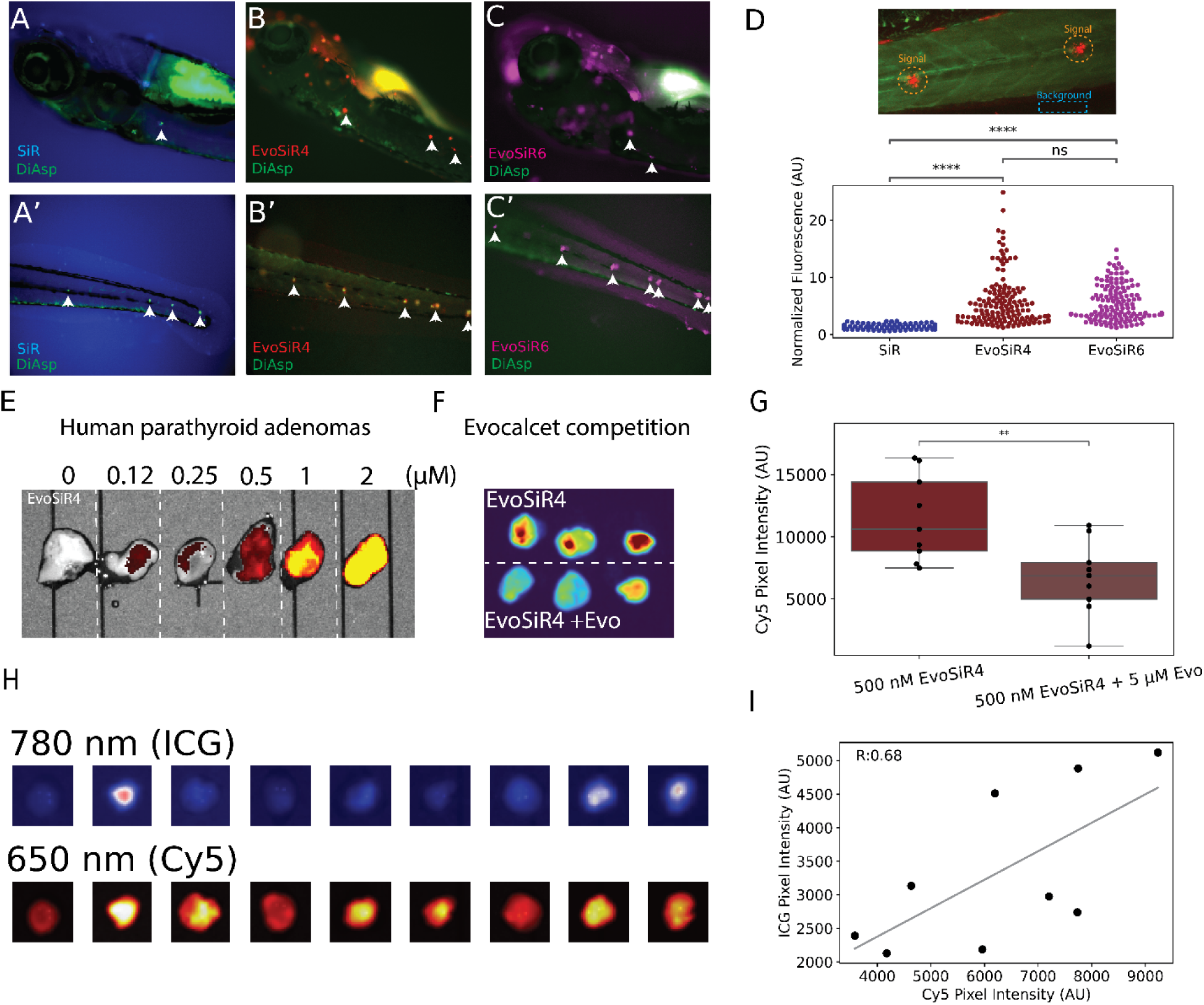
EvoSiR probes specifically label neuromasts in zebrafish larvae. (A-C) Representative images of live zebrafish larvae co-stained with SiR (**4**) or EvoSiR probes and the specific neuromast label, DiAsp. (D) The top image shows the EvoSiR4 labelling of neuromasts acquired by confocal microscopy. The bottom plot shows the normalized fluorescence of the neuromasts identified with the DiAsp staining to the background fluorescence value for each probe n =14-18; p=0.64 for comparing EvoSiR4 and EvoSiR6, p<0.0001 for all other comparisons, Mann-Whitney Wilcoxon test with Bonferroni correction). (E) Concentration-dependence of EvoSiR4 labelling on *ex vivo* human parathyroid adenomas. At 650 nm excitation, no autofluorescence of the tissue is observed, whereas the addition of >500 nM of EvoSiR4 results in a marked increase in tissue fluorescence. (F) Representative images of three parathyroid adenomas cut in half, where the top halves were labelled with 500 nM of EvoSiR4, whereas the bottom halves co-administered with 500 nM of EvoSiR4 and 2.5µM of evocalcet. A significant decrease in tissue fluorescence was observed upon evocalcet competition. (G) Boxplot depicting the results of evocalcet competition experiments on nine parathyroid adenomas (p=0.006, n=9 each, Student’s t-test). (H) Representative images of the autofluorescence at 780 nm (ICG channel) and EvoSiR at 650 nm (Cy5 channel) of nine labelled, surgically resected ex vivo human parathyroid adenomas. I: Plot depicting a significant positive correlation between the average pixel intensity of the Cy5 channel (EvoSiR) and the ICG channel (autofluorescence) (n=9, Pearson’s R=0.676, p=0.04).

### EvoSiR had no effect on the modulation of startle reflex in zebrafish larvae

To investigate whether EvoSiR can penetrate into the brain of zebrafish larvae, we quantified the latency of the startle reflex in response to acoustic-vibrational stimuli, which has been previously reported to be modulated by CaSR^10,38^. In response to abrupt low-level acoustic stimulation, zebrafish elicited the startle reflex which comprised of short-latency responses (2-12 ms) in 12%, long-latency responses (>12 ms) in 3%, and no response in 85% of the total number of stimulations. Bath-applied EvoSiR4 appeared to have no effect on the basic locomotion capabilities of the larvae at up to 10 µM, however it was lethal beyond 20 µM. We found no significant difference in neither the percentage of elicited startle responses, nor the ratio of short-latency escapes (Figure S4). To assess target engagement of EvoSiR4 in the brain, we imaged the transgenic *Tol056* zebrafish strain, which express eGFP in the Mauthner-neurons in the hindbrain responsible for the initiation of the short-latency escape response using confocal microscopy^36^. In agreement with the behavioral data, we found no significant EvoSiR label, indicating that the bath-applied probe does not enter the zebrafish CNS (Figure S4).

### EvoSiR labelling of ex vivo human parathyroid adenomas

To assess the utility of EvoSiR in labeling CaSR expressing human tissues, we applied EvoSiR4 to ex vivo surgically resected human parathyroid adenomas, which showed a dose-dependent increase in tissue fluorescence at 650 nm upon the addition of EvoSiR4 (Figure 5). We next conducted competition experiments in which tissues were halved and one half was co-incubated with EvoSiR4 and evocalcet. We observed a significant decrease in the absolute fluorescence in the evocalcet-competed tissues (Figure 5). Next, we stained nine additional adenomas and compared the labeling intensity with the autofluorescence of the tissues at 780 nm. The intensity of near infrared (NIR) autofluorescence and CaSR expression of the parathyroid glands have been shown to be significantly lower in adenomas compared to the normal parathyroid glands^39^. We acquired images at 780 nm and 650 nm in parallel, which showed a significant positive correlation (r = 0.68) between the autofluorescence and the EvoSiR4 labeling, which might indicate that EvoSiR4 remains specific towards CaSR in human tissues (Figure 5).

## Discussion

Mounting transdisciplinary research interest on the interrogation of CaSR signaling warrants the development of fast, broadly applicable and versatile molecular tools for its visualization and the perturbation of its function. In the current study, we describe the design, synthesis, and biological characterization of novel SiR functionalized CaSR targeting fluorescent probes of evocalcet called EvoSiR. The *in silico* and *in vitro* data show that evocalcet is tolerant towards the introduction of large chemical tags at the carboxyl end, which renders the potently CaSR interacting chemical scaffold optimal for the development of chemical probes. Using spectrophotometric approaches, we demonstrate that EvoSiR probes are not only potent CaSR ligands, but also efficiently bind BSA, enabling the removal of non-specific signals. This was supported by efficient *in vitro* labeling and competition experiments on a CaSR overexpressing cell line. Furthermore, we confirmed the *in vivo* applicability of EvoSiR4 and EvoSiR6, which specifically labelled the hair cells in live zebrafish larvae, Lastly, we explored the utility of EvoSiR in labeling *ex vivo* human parathyroid adenomas without BSA washing, showing a strong correlation with parathyroid glands, and a two-fold increase to the autofluorescence signal.

From the diverse small molecules that have been developed to target CaSR^18^, the selection of a chemical scaffold suitable for the incorporation of fluorescent tags could be ambiguous. Indeed, our initial strategy of linking an NBD fluorophore to the *meta*-position of the *N*-benzyl group of quinazoline-2-ones, similar to the potent calcilytic compounds ATF936 and AXT914^17^, resulted in diminished potency towards CaSR, in agreement with previously published SAR data^40^, rendering the molecule inadequate for CaSR-specific labeling purposes (data not shown). In addition, these *N*-benzyl quinazoline-2-ones are very lipophilic compounds per se that ultimately resulted in considerable non-specific labelling of the fluorescent NBD-conjugates. In contrast, SAR studies on the evocalcet structure provided evidence that the carboxyl position of the molecule is tolerant towards structural modifications. In agreement, recent Cryo-EM studies have shown that in the CaSR-bound state, the carboxyl position of evocalcet is less constrained within the neighboring environment^21,22^. To acquire a comprehensive picture of potentially suitable chemical scaffolds, we evaluated all quantified structures from SAR studies available from PubChem. In accordance with the literature data on evocalcet, the analysis revealed this structure to be the least sensitive to structural modifications, which led us to investigate the impact of fluorophore conjugation on its potency and specificity on CaSR.

Using a functional CaSR assay and *in vitro* labeling experiments, we showed that EvoSiR improves with a 6-fold increase on CaSR activity compared to evocalcet (EC_50_ = 243 nM), with an EC_50_ value in the low nanomolar ranges (<50 nM) and remains specific towards the receptor below 500 nM. As shown in a previous report in which the SAR of evocalcet structures was evaluated, several analogues modified at the carboxyl position exhibited higher CaSR activity, whereas evocalcet had the most favorable PK profile with no direct CYP2D6 inhibition^41^. Of note, the four-carbon linker conjugation to evocalcet (compound **1**, Scheme 1) also reduced the EC_50_ value to 101 nM. Therefore, it is plausible that the further optimization of the carboxyl position of evocalcet could yield molecules with even higher activity in the future.

Using spectrofluorometry, we revealed the strong BSA binding of EvoSiR, which was consistent with the observed concentration-dependent decrease in CaSR affinity upon the addition of BSA in the functional assay. Albumin binding is a critical parameter to optimize in the design of *in vivo* experiments and potential clinical translation and was different for the two EvoSiR probes EvoSiR4 and EvoSiR6. Two experimental results support the potential positive effect of BSA on EvoSiR target engagement. First, the CaSR specific labelling of cells resulted in an increased signal-to-background ratio. Second, our *in vivo* data on labeling the neuromasts of zebrafish larvae showed a superior signal-to-noise ratio of EvoSiR to existing labels (DiAsp), which suggests that the high albumin binding of the molecule may improve its labeling capacity. Still, it remains to be elucidated in a rodent model and in humans whether such high albumin affinity would result in diminished target engagement; or would improve the signal-to-noise ratio by washing out non-specific weak binding interactions from the target site.

In the removal of solid tumors and the preservation of healthy tissues during surgery, the application of fluorescent dyes such as Indocyanine Green (ICG)^42^ and the protoporphyrin IX precursor 5-aminolevulinic acid (5-ALA)^43^ has notable clinical utility. However, the feasibility of applying always-on fluorescent probes in the fluorescence-guided surgery context, especially in the form of a topical sprays can be impaired by low target specificity and high background fluorescence^44^. Activatable fluorescent probes such as enzyme-cleavable probes or pH-activable probes showed promising results in the accurate determination of tumor margins *in situ*^45,46^. Moreover, recent developments of multivariate AND-gate optical contrast agents, which produce a signal upon least at two proteolytic processes may further enhance the contrast between the tumor boundaries and surrounding healthy tissues^47^.

On the translational perspective, the visualization of CaSR in the human parathyroid glands could enhance the localization and thus preservation of the tissue, which is an integral objective during various head and neck surgical procedures. For example, during thyroidectomy, which is performed approximately 100,000 times annually in just the US, transient and persistent damage to the parathyroid glands represent a major complication in up to 30% and 3% of the cases, respectively^48^. Currently the relatively high far-red autofluorescence of the parathyroid glands compared to the surrounding tissues is used for parathyroid gland identification, which does not inform surgeons about the perfusion and the vitality of the glands during the operation^49^. For parathyroid gland perfusion, indocyanine green angiography can be applied, which is an aspecific label^50^. Altough the recently developed structure-inherent probe, T800-F that is specifically taken up by the parathyroid glands is chemically more compact, its flexibility to optimize for clinical use (DMPK, toxicity) might be limited^51^. The application of small molecule CaSR-specific probes like EvoSiR4 for fluorescence-guided thyroidectomy could therefore help in both PG identification and assessing PG perfusion of the surrounding vasculature. EvoSiR probes exhibit nanomolar potency towards CaSR, ensuring specificity. Moreover, the probes possess a high affinity for albumin, a characteristic that may facilitate the removal of EvoSiR not bound to CaSR during the washing process. In addition, the fluorogenicity of the probe upon albumin binding may enable the visualization of the vasculature of the PG, allowing the assessment of both the anatomy and the perfusion parameter with a single probe.

Our results also support the possibility of applying EvoSiR probes as an intraoperative topical spray for parathyroid gland identification *in vivo* in the future, because the highly potent SiR quencher tannic acid, identified in this study, may increase target-related fluorescence by selectively quenching only the EvoSiR not bound to CaSR. Our *in vitro* data supports this possibility, as the addition of tannic acid significantly improved the signal to background ratio in cellular CaSR labelling experiments. Although an *in vivo* applicability of this technique may be challenging, EvoSiR probes have translational potential for the development of optimized probes that can inform surgeons on the anatomical identification of CaSR expressing parathyroid glands in excised tissues^52^. Therefore, considering ADMETOX properties, the labelling capacity of EvoSiR probes will be explored in preclinical *in vivo* models.

### Conclusions

To our knowledge, the SiR conjugates of the carboxyl end of evocalcet (EvoSiR) are the first fluorescent CaSR probes which show low nanomolar CaSR activity and improved potency over evocalcet. The CaSR labelling capacity of EvoSiR probes were validated in various *in vitro*, *in vivo* settings and *ex vivo* human parathyroid tissues. The tolerance of evocalcet towards functionalization with fluorophores or other tags permits large-scale optimization for biomedical applications.

## Author contributions

Conceptualization, JG, RMK, ML and DB; Chemical synthesis, JPF; Methodology, DB, JPF; Programming and data analysis, DB; Visualization, DB, ML; Manuscript writing (draft preparation), DB, JG; Manuscript writing (review and editing), all authors.

## Acknowledgements

We would like to thank XY from Novartis for kindly providing us the calcium-sensing receptor expressing cell line. We would like to further thank Prof. András Málnási-Csizmadia and his lab (Department of Biochemistry, ELTE Eötvös Loránd University, Budapest, Hungary) for their kind help with the *in vivo* experiments and acknowledge Dr. Katalin Schlett and Dr. Norbert Bencsik (Department of Physiology, ELTE Eötvös Loránd University, Budapest, Hungary) for their help with setting up the confocal microscopy experiments. We thank the Analytical Services from the Department of Chemistry, Biochemistry, and Pharmaceutical Sciences (DCBP), University of Bern, Switzerland (Prof. Julien Furrer; Prof. Stefan Schürch), for measuring NMR and MS spectra of synthetic intermediates and final compounds.

