## Supporting Information for "Silicon-rhodamine functionalized evocalcet probes (EvoSiR) potently and selectively label calcium sensing receptors (CaSR) *in vitro*, *in vivo* and *ex vivo*"

Click or tap here to enter text.

### Supporting Information

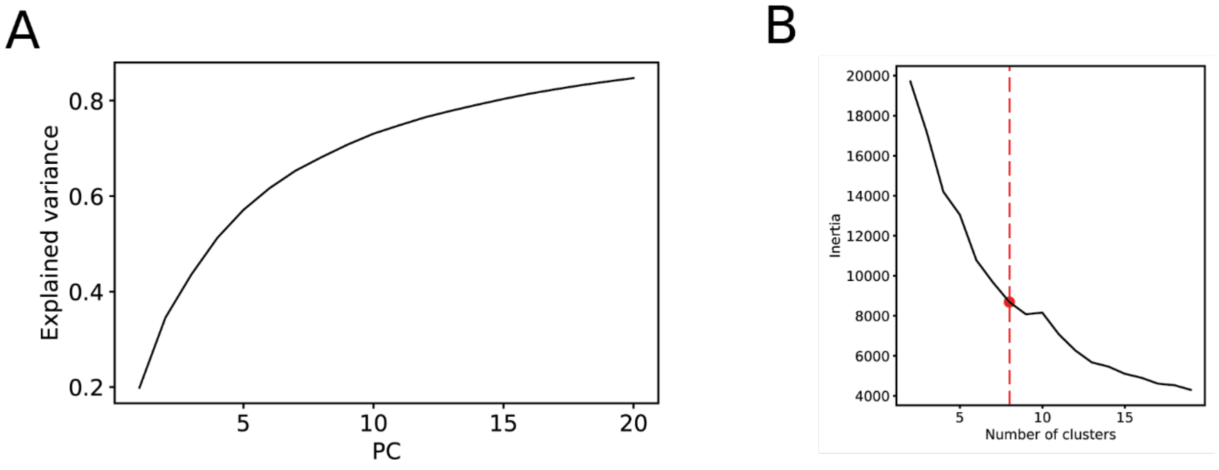


**Figure S1. Properties of the PCA and k-means algorithms.** (A) Cumulative explained variance of 20 principal components (PC) conducted on the PubChem fingerprints. (B) Selection of the optimal number of clusters based on the inertia score

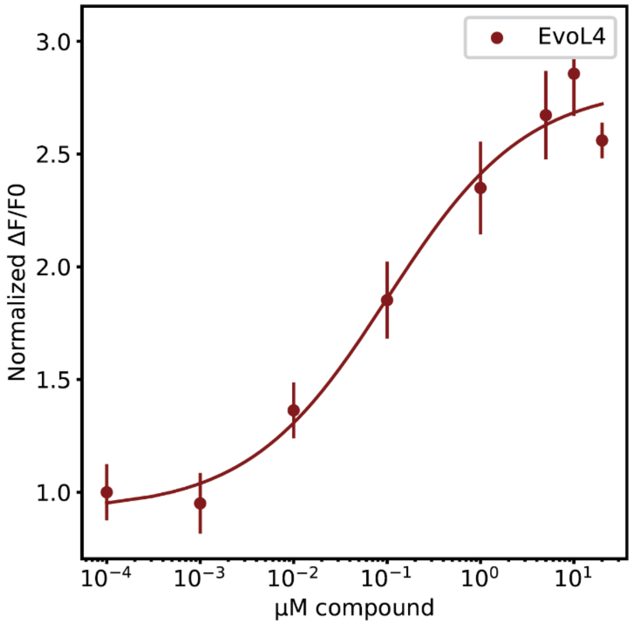


**Figure S2. Dose-response curve of EvoL4 (1).** FLIPR assay measuring intracellular calcium levels of evocalcet conjugated to the four-carbon linker (n=4, in triplicates, $EC_{50}$=0.101±0.006 μM)

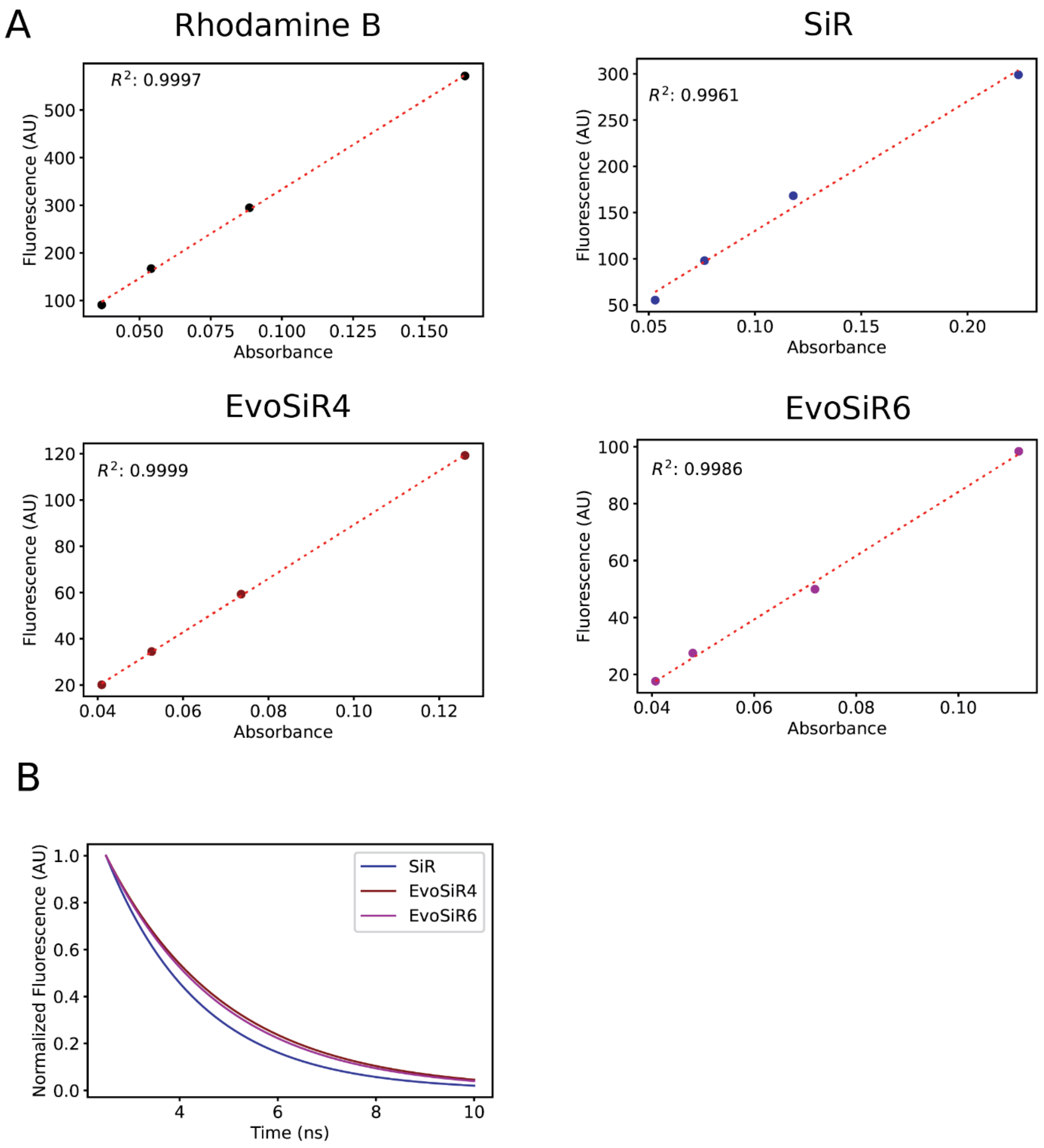


**Figure S3. Quantum yield and fluorescence lifetime measurements.** (A) Linear regression curves used for the estimation of relative quantum-yield for the reference compound (Rhod B) and the SiR (**4**) and EvoSiR probes (n=3 each). (B) Exponential fits for the three compounds used for the estimation of fluorescence lifetimes.

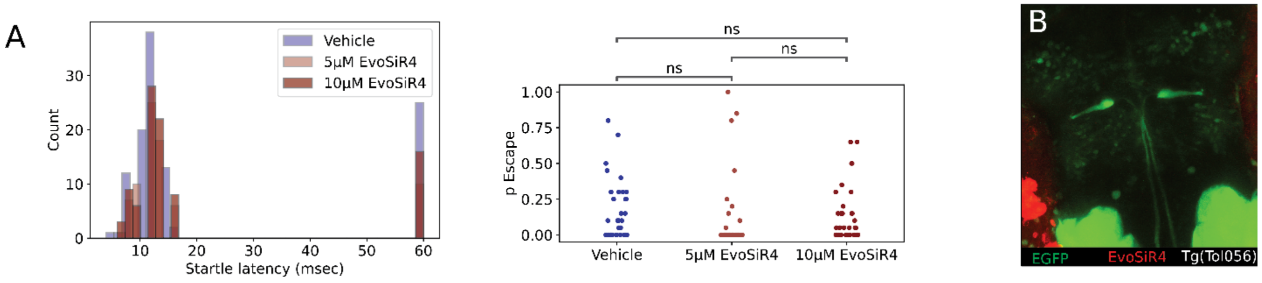


**Figure S4. The effect of EvoSiR4 on zebrafish startle behavior** (A) The latency of startle responses for n=36 fish per condition shows no difference between the vehicle controls and the bath administration EvoSiR4 (left panel). The probability of eliciting an escape response does not significantly change upon the administration of EvoSiR4 (n = 36 fish per condition, p =0.06, 0.1, 0.99 for the comparison of vehicle and 5 µM EvoSiR4, vehicle and 10 µM EvoSiR4 and 5 µM EvoSiR4 and 10 µM EvoSiR4, respectively: Mann-Whitney Wilcoxon test with Bonferroni correction). (B) Image of the Mauthner-neuron labelled with 5µM EvoSiR4. No EvoSiR4 signal was detected inside the brain, whereas the superficial layers were labelled.

**Synthesis of chemical compounds**

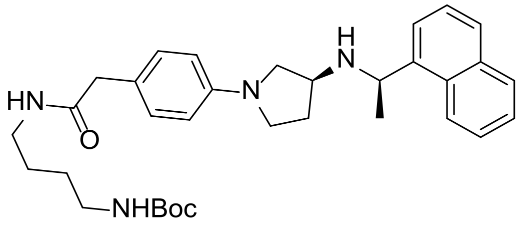


**Tert-butyl (4-(2-(4-((S)-3-(((R)-1-(naphthalen-1-yl)ethyl)amino) pyrrolidin-1-yl)phenyl) acetamido) butyl) carbamate (1)**

To a solution of Evocalcet (0.13 mmol, 50 mg, 1 eq) in dichloromethane (5 mL) was added N-(3-dimethylaminopropyl)-N′-ethylcarbodiimide hydrochloride (0.26 mmol, 50 mg, 2 eq) at 0°C and the suspension was stirred at this temperature for 30 min. Then, 1-Boc-1,4-butanediamine (0.15 mmol, 28 µL, 1.1 eq) and 4-(dimethylamino) pyridine (0.16 mmol, 19 mg, 1.2 eq) were added at 0°C and the reaction mixture was stirred at RT for 16h. Water (5 mL) was added, and the product was extracted with EtOAc (3x5 mL). The organic phase was dried over sodium sulphate and the solvent was removed in vacuo. The crude was purified by reverse-phase flash-chromatography (water + 0.1%TFA/acetonitrile + 0.1%TFA) to obtain the desired product (61 mg, 84%).

**$^{1}$H NMR** (300 MHz, $CDCl_3$) δ 10.98 – 9.64 (m, 3H), 8.11 – 7.85 (m, 4H), 7.74 – 7.49 (m, 3H), 6.96 (d, J = 7.9 Hz, 2H), 6.35 (d, J = 8.0 Hz, 2H), 5.45 – 5.30 (m, 1H), 3.66 – 3.27 (m, 3H), 3.39 (s, 2H), 3.27 – 3.17 (m, 1H), 3.17 – 3.07 (m, 2H), 3.07 – 2.92 (m, 3H), 2.53 – 2.31 (m, 1H), 2.20 – 2.04 (m, 1H), 1.78 (d, J = 6.4 Hz, 3H), 1.52 – 1.30 (m, 4H), 1.42 (s, 9H).

**$^{13}$C NMR** (75 MHz, $CDCl_3$) δ 145.92, 141.19, 136.02, 134.01, 131.85, 130.62, 130.35, 129.94, 129.63, 127.64, 126.50, 125.89, 124.86, 122.72, 121.10, 117.48, 112.93, 99.15, 92.41, 90.82, 55.19, 50.67, 42.09, 39.48, 28.37, 27.62, 26.14, 24.23, 21.01.

**FTMS** (NSI +) Calculated for $C_{33}H_{45}N_4O_3$ [M–H]+: 545.3486 found: 545.3475.

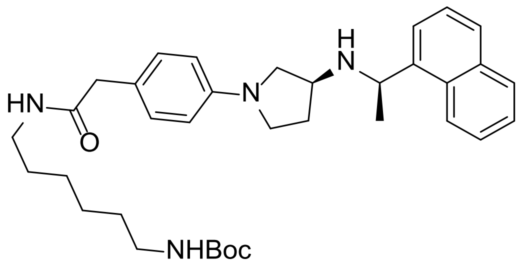


**Tert-butyl (6-(2-(4-((S)-3-(((R)-1-(naphthalen-1-yl)ethyl)amino) pyrrolidin-1-yl)phenyl) acetamido) hexyl) carbamate (2)**

To a solution of evocalcet (0.13 mmol, 50 mg, 1 eq) in dichloromethane (4 mL) was added N-(3-dimethylaminopropyl)-N′-ethylcarbodiimide hydrochloride (0.26 mmol, 50 mg, 2 eq) at 0°C and the suspension was stirred at this temperature for 30 min. Then, a solution of 1-Boc-1,6-hexanediamine (0.18 mmol, 40 mg, 1.4 eq) and 4-(dimethylamino) pyridine (0.16 mmol, 19 mg, 1.2 eq) in dichloromethane (1 mL) was added at 0°C and the reaction mixture was stirred at RT for 16h. Water (5 mL) was added, and the product was extracted with EtOAc (3x5 mL). The organic phase was dried over sodium sulphate and the solvent was removed in vacuo. The crude was first purified by reverse-phase flash-chromatography (water + 0.1%TFA/acetonitrile + 0.1%TFA) and then by normal phase (dichloromethane/methanol) to obtain the desired product (47 mg, 62%).

**$^{1}$H NMR** (300 MHz, $CDCl_3$) δ 8.27 – 8.17 (m, 1H), 7.93 – 7.83 (m, 1H), 7.81 – 7.64 (m, 2H), 7.56 – 7.41 (m, 3H), 7.04 (d, 2H), 6.45 (d, 2H), 5.50 (s, 1H), 4.80 (q, J = 6.5 Hz, 1H), 4.63 (s, 1H), 3.46 – 3.34 (m, 3H), 3.44 (s, 2H), 3.24 – 3.10 (m, 3H), 3.10 – 2.99 (m, 3H), 2.25 – 2.11 (m, 1H), 1.98 – 1.78 (m, 2H), 1.53 (d, J = 6.5 Hz, 3H), 1.47 – 1.31 (m, 4H), 1.44 (s, 9H), 1.28 – 1.18 (m, 4H).

**$^{13}$C NMR** (75 MHz, $CDCl_3$) δ 172.20, 156.06, 147.01, 140.88, 134.04, 131.28, 130.33, 129.09, 127.42, 125.91, 125.71, 125.43, 123.18, 122.83, 121.48, 111.98, 79.00, 55.51, 54.11, 51.79, 46.30, 42.97, 40.41, 39.38, 32.02, 29.95, 29.41, 28.49, 26.36, 26.29, 24.17.

**FTMS** (NSI +) Calculated for $C_{35}H_{49}N_4O_3$ [M–H]+: 573.3799 found: 573.3808.

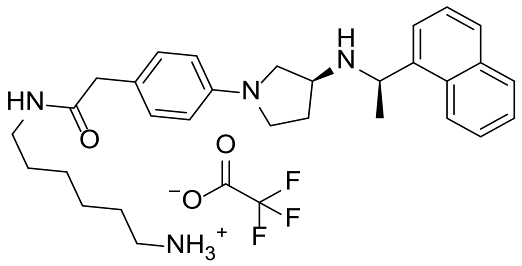


**6-(2-(4-((S)-3-(((R)-1-(naphthalen-1-yl)ethyl)amino) pyrrolidin-1-yl) phenyl) acetamido) hexan-1-aminium 2,2,2-trifluoroacetate (3)**

To a solution of **2** (0.117 mmol, 67 mg, 1 eq) in dichloromethane (3 mL) was added trifluoroacetic acid (1.298 mmol, 0.1 mL, 11.1 eq) and the reaction mixture was stirred for 2h at room temperature. The reaction mixture was concentrated in vacuum without heating, the residual oil was dissolved in dichloromethane, and concentrated again. The crude was purified by flash chromatography (dichloromethane / methanol) to obtain the title compound (61 mg, 89%).

**$^{1}$H NMR** (300 MHz, MeOD) δ 8.23 (d, *J* = 8.5 Hz, 1H), 8.01 – 7.87 (m, 2H), 7.81 (dd, *J* = 7.3, 1.2 Hz, 1H), 7.68 – 7.46 (m, 3H), 7.05 (d, *J* = 8.6 Hz, 2H), 6.43 (d, *J* = 8.7 Hz, 2H), 5.48 (q, *J* = 6.7 Hz, 1H), 3.84 – 3.72 (m, 1H), 3.49 – 3.38 (m, 1H), 3.32 (s, 2H), 3.30 – 3.25 (m, 2H), 3.16 – 2.99 (m, 3H), 2.84 (t, *J* = 7.6 Hz, 2H), 2.27 (q, *J* = 7.0 Hz, 2H), 1.81 (d, *J* = 6.6 Hz, 3H), 1.65 – 1.52 (m, 2H), 1.52 – 1.40 (m, 2H), 1.35 – 1.22 (m, 4H).

**$^{13}$C NMR** (75 MHz, MeOD) δ 174.78, 163.24 (TFA), 162.79 (TFA), 147.62, 135.51, 133.95, 132.02, 131.06, 130.74, 130.43, 128.62, 127.59, 126.68, 125.51, 125.42, 122.85, 120.15 (TFA), 116.27 (TFA), 113.71, 56.60, 53.20, 51.54, 46.93, 42.99, 40.53, 40.19, 30.05, 29.34, 28.35, 27.20, 26.89, 20.55.

**FTMS** (NSI +) Calculated for $C_{30}H_{41}N_4O$ [M–H]+: 473.3275 found: 473.3262.

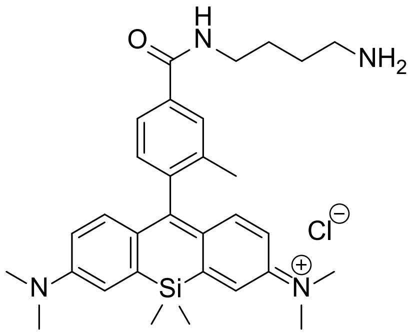


**N-(10-(4-((4-aminobutyl) carbamoyl)-2-methylphenyl)-7-(dimethylamino)-5,5-dimethyldibenzo [b,e] silin-3 (5H)-ylidene)-N-methylmethanaminium chloride (4, SiR)**

To a solution of **SiR-$CO_2$H** (0.067 mmol, 40 mg, 1 eq) in dichloromethane (5 mL) was added N-(3-dimethylaminopropyl)-N′-ethylcarbodiimide hydrochloride (0.14 mmol, 27 mg, 2 eq) at 0°C and the suspension was stirred at this temperature for 30 min. Then, 1-Boc-1,4-butanediamine (0.078 mmol, 15 mg, 1.1 eq) and DMAP (0.0819 mmol, 10 mg, 1.2 eq) were added at 0°C and the reaction mixture was stirred at room temperature for 16h. Then, water (5 mL) was added, and the product was extracted with EtOAc (3x5 mL). The organic phase was dried over sodium sulphate and concentrated in vacuo. The crude was purified by flash liquid chromatography (methanol/dichloromethane) to give the conjugated product.

This product was dissolved in acetonitrile (1 mL) and aq. HCl (6 M, 4 mL) was added. The blue solution was stirred at RT for 2 h. The solution was cooled to 0°C and aq. NaOH (2 M, 5.4 mL) was slowly added. The resulting mixture was then concentrated in vacuo to give a crude product, in presence of NaCl. This was dissolved in methanol, filtrated, and purified by flash chromatography (DCM/MeOH) to give the desired product (42 mg, quant. over 2 steps).

**$^1$H NMR** (300 MHz, MeOD) δ 7.92 (d, *J* = 1.8 Hz, 1H), 7.87 (dd, *J* = 7.9, 1.8 Hz, 1H), 7.38 (d, *J* = 2.8 Hz, 2H), 7.26 (d, *J* = 7.9 Hz, 1H), 7.03 (d, *J* = 9.6 Hz, 2H), 6.78 (dd, *J* = 9.7, 2.9 Hz, 2H), 3.55 – 3.44 (m, 2H), 3.35 (d, *J* = 2.7 Hz, 13H), 3.06 – 2.99 (m, 2H), 2.11 (s, 3H), 1.85 – 1.71 (m, 4H), 0.62 (d, *J* = 4.5 Hz, 6H).

**$^{13}$C NMR** (75 MHz, MeOD) δ 169.51, 155.81, 149.49, 141.91, 137.66, 136.09, 130.53, 130.22, 128.07, 125.81, 122.37, 115.37, 68.24, 41.01, 40.46, 27.48, 25.97, 19.50, -1.06, -1.28.

**FTMS** (NSI +) Calculated for $C_{31}H_{41}N_4OSi$ [M]+: 513.3044 found: 513.3039.

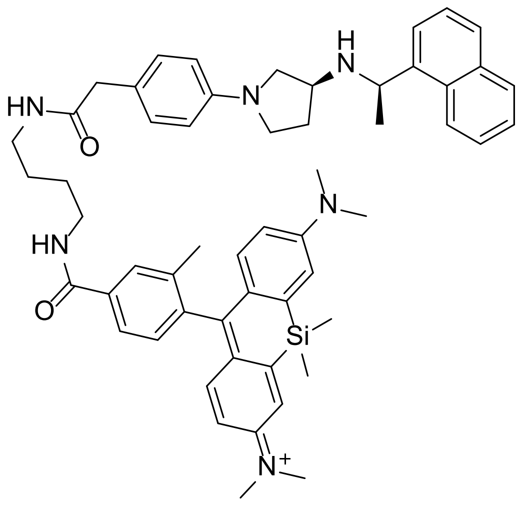


**N-(7-(dimethylamino)-5,5-dimethyl-10- (2-methyl-4-((4-(2-(4-((S)-3-(((R)-1-(naphthalen-1-yl) ethyl) amino) pyrrolidin-1-yl) phenyl) acetamido) butyl) carbamoyl) phenyl) dibenzo[b,e]silin-3(5H)-ylidene)-N-methylmethanaminium (EvoSiR4)**

To a solution of evocalcet (0.069 mmol, 26 mg, 1 eq) in dichloromethane (2 mL) was added N-(3-dimethylaminopropyl)-N′-ethylcarbodiimide hydrochloride (0.136 mmol, 26 mg, 2 eq) at 0°C and the suspension was stirred at this temperature for 30 min. Then, **X4** (0.069 mmol, 40 mg, 1 eq) in DCM (5 mL) and DMAP (0.164 mmol, 20 mg, 2.4 eq) were added at 0°C and the reaction mixture was stirred at RT for 16h. The mixture was concentrated in vacuo, and the crude was purified by flash chromatography (dichloromethane/methanol) and reverse-phase flash chromatography (water + 0.1%TFA/acetonitrile + 0.1%TFA) to give the desired product (9 mg, 14%). Purity (HPLC): >99.9% (detection at 254 nm).

**$^{1}$H NMR** (300 MHz, MeOD) δ 8.26 (d, *J* = 8.4 Hz, 1H), 8.06 – 7.96 (m, 2H), 7.90 – 7.75 (m, 3H), 7.70 – 7.55 (m, 3H), 7.37 (d, *J* = 2.8 Hz, 2H), 7.23 (d, *J* = 7.8 Hz, 1H), 7.13 (d, *J* = 8.5 Hz, 2H), 7.03 (d, *J* = 9.7 Hz, 2H), 6.76 (dd, *J* = 9.7, 2.9 Hz, 2H), 6.54 (d, *J* = 8.6 Hz, 2H), 5.53 (q, *J* = 6.7 Hz, 1H), 3.92 (t, *J* = 6.4 Hz, 1H), 3.53 (td, *J* = 9.1, 4.7 Hz, 1H), 3.46 – 3.33 (m, 6H), 3.35 (s, 12H), 3.27 – 3.14 (m, 3H), 2.43 – 2.24 (m, 2H), 2.09 (s, 3H), 1.86 (d, *J* = 6.7 Hz, 3H), 1.70 – 1.55 (m, 4H), 0.61 (d, *J* = 4.6 Hz, 6H).

**$^{13}$C NMR** (101 MHz, MeOD) δ 173.47, 167.99, 167.87, 154.43, 148.11, 146.30, 142.00, 140.55, 136.24, 134.94, 134.21, 132.56, 130.65, 129.84, 129.42, 129.11, 128.73, 127.31, 126.70, 126.30, 125.28, 124.31, 124.28, 121.44, 120.93, 113.93, 112.48, 55.38, 50.24, 47.60, 45.63, 41.66, 39.52, 39.28, 38.82, 28.16, 26.53, 26.47, 18.88, 18.10, -2.51, -2.73.

**FTMS** (NSI +) Calculated for $C_{55}H_{65}N_{6}O_{2}Si$ [M]+: 869.4933 found: 869.4926.

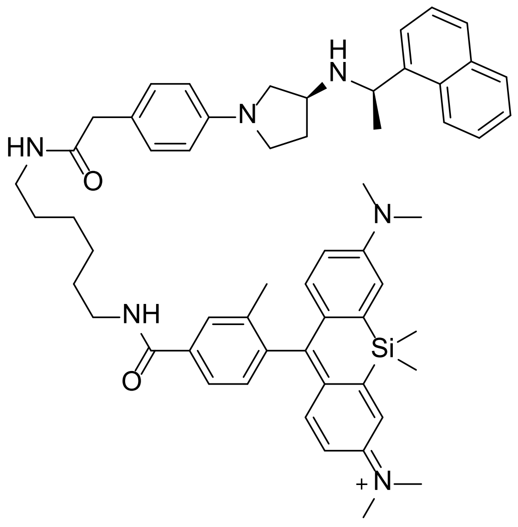

**N-(7-(dimethylamino)-5,5-dimethyl-10- (2-methyl-4-((6-(2-(4-((S)-3-(((R)-1-(naphthalen-1-yl) ethyl) amino) pyrrolidin-1-yl) phenyl) acetamido) hexyl) carbamoyl) phenyl) dibenzo[b,e]silin-3(5H)-ylidene)-N-methylmethanaminium (EvoSiR6)**

To a solution of **SiR-NHS** (0.023 mmol, 13 mg, 0.8 eq) in acetonitrile (1 mL) was added **3** (0.029 mmol, 17 mg, 1 eq) in acetonitrile (2 mL) and DIPEA (0.075 mmol, 13 µL, 2.6 eq). The mixture was stirred for 20h at room temperature and concentrated in vacuum. The crude was purified by flash chromatography (dichloromethane/methanol) and reverse-phase chromatography (water + 0.1%TFA/acetonitrile + 0.1%TFA) to give the desired product (7 mg, 25%). Purity (HPLC): >95% (detection at 254 nm).

**$^{1}$H NMR** (300 MHz, MeOD) δ 8.26 (d, *J* = 8.4 Hz, 1H), 8.04 – 7.95 (m, 2H), 7.89 – 7.77 (m, 3H), 7.69 – 7.54 (m, 3H), 7.37 (d, *J* = 2.9 Hz, 2H), 7.21 (d, *J* = 7.9 Hz, 1H), 7.12 (d, *J* = 8.3 Hz, 2H), 7.02 (d, *J* = 9.7 Hz, 2H), 6.74 (dd, *J* = 9.6, 2.8 Hz, 2H), 6.57 – 6.48 (m, 2H), 5.53 (q, *J* = 6.7 Hz, 1H), 3.91 (q, *J* = 6.4 Hz, 1H), 3.52 (td, *J* = 8.6, 4.6 Hz, 1H), 3.42 – 3.36 (m, 4H), 3.35 (s, 12H), 3.17 (t, *J* = 7.0 Hz, 3H), 2.46 – 2.22 (m, 2H), 2.08 (s, 3H), 1.85 (d, *J* = 6.7 Hz, 3H), 1.69 – 1.57 (m, 2H), 1.57 – 1.46 (m, 2H), 1.46 – 1.23 (m, 6H), 0.61 (d, *J* = 5.1 Hz, 6H).

**FTMS** (NSI +) Calculated for $C_{57}H_{69}N_6O_2Si$ [M]+: 897.5246 found: 897.5258.
